# A biphasic Deep Semi-supervised framework for Suptype Classification and biomarker discovery

**DOI:** 10.1101/2022.01.13.476268

**Authors:** Hoang Le, Van-Minh Nguyen, Quang-Huy Nguyen, Duc-Hau Le

## Abstract

To take full advantage of the unprecedented development of -omics technologies and generate further biological insights into human disease, it is a pressing need to develop novel computational methods for integrative analysis of multi-omics data. Here we proposed a biphasic Deep Semi-supervised multi-omics integration framework for Subtype Classification and biomarker discovery, DeepSSC. In phase 1, each denoising autoencoder was used to extract a compact representation for each -omics data, and then they were concatenated and put into a feed-forward neural network for subtype classification. In phase 2, our Biomarker Gene Identification procedure leveraged that neural network classifier to render subtype-specific important biomarkers. We also validated our given results on independent dataset. We demonstrated that DeepSSC exhibited better performance over other state-of-the-art techniques concerning classification tasks. As a result, DeepSSC successfully detected well-known biomarkers and hinted at novel candidates from different -omics data types related to the investigated biomedical problems.

## INTRODUCTION

Proliferation of various types of “-omics” data with unprecedented details until now has been caused by the rapid development in high-throughput biomedical technologies. This has enabled us to build more complicated models by combining multiple -omics (multi-omics) for various studies of diseases. It is common to encounter the claim that integrating multiple types of omics data help us to discover a more holistic view of the underlying biological processes rather than when using each omics technology [1–3]. Therefore, there is a pressing need for us to build novel integrative analysis methods that may effectively leverage the complementary information in multi-omics data.

Over the past years, cancer subtypes have been established gradually and inheritably to hope that we can build individualized treatment strategies for cancer patients. Currently, there are two key problems researchers strive to solve. On the one hand, a lot of literature are driven in the direction of classifying tumors using various -omics data types such as miRNA arrays [4, 5], copy-number alterations [6] or an integration of different -omics datasets [3, 7, 8]. On the other hand, many works seek to re-cluster cancer patients in the hope that they may find out novel patient subgroups [3, 9]. Supervised and unsupervised machine learning techniques are used widely to rectify these problems. Then, deep learning has been leveraged to change the way the data are processed as the discovery of features are done rapidly with little human intervention or domain knowledge. Deep neural networks help develop predictive models that deal efficiently with the challenges of high-dimensional data [10]. Indeed, high-end deep learning frameworks often outperform machine learning approaches with regard to computational performance [11, 12], such as breast cancer (BRCA) classification using genomic data [13, 14] or gene expression inference [15, 16] or Alzheimer’s disease classification using integrative -omics data [17].

Numerous computational methods established at first for classification problems have focused on unsupervised multi-omics integration without the additional information of sample labels, such as [7, 18]. With the unprecedented proliferation of omics technologies and the establishment of cancer genomic databases, the availability of labeled omics datasets characterizing the phenotypes or clinical features of the samples are becoming more widely common. Therefore, today researchers have been of increasing interest to problems associated with disease-related biomarker identification and predictions on novel samples by adopting supervised multi-omics integration approaches, such as [17, 19]. However, to the best of our knowledge, it is very rare to see or even has never ever seen semi-supervised learning used for -omics-based classification tasks in biomedical applications, although it is considered as a merger of the strengths of the supervised and unsupervised learning.

In this study, we introduce a novel biphasic Deep Semi-supervised multi-omics integration framework, called DeepSSC, that can be able to implement two tasks: (i) cancer subtype classification and (ii) subtype-specific biomarker identification through a range of biomedical classification applications, including BRCA, colon & rectum adenocarcinoma (COADREAD), and lung cancer. We also open the new way for applying our semi-supervised learning strategy to biomedical classification problems of -omics data and show its effectiveness as well. To be specific, we employ two -omics profiles, including mRNA expression (GE) and copy-number alterations (CNA), downloaded from two independent sources The Cancer Genome Atlas (TCGA) [20] and the Molecular Taxonomy of Breast Cancer International Consortium (METABRIC) [21] (Figure 1). The first phase uses separately each denoising autoencoder (DAE) for each -omics profile, rendering two respective latent-space representations. These representations are then concatenated and transformed through two fully-connected (FC) layers to build a neural network classifier for cancer subtype classification. At last, we propose a Biomarker Gene Identification (BGI) procedure in the second phase to identify subtype-specific important biomarkers with the input being the above trained neural network classifier. We will show that our identified biomarkers are well able to help classify cancer patients into different subtypes even if only using traditional machine learning models in phase 2. We also further demonstrate that DeepSSC outperforms other state-of-the-art supervised multi-omics data classification methods regarding cancer subtype classification.

**Figure 1.**
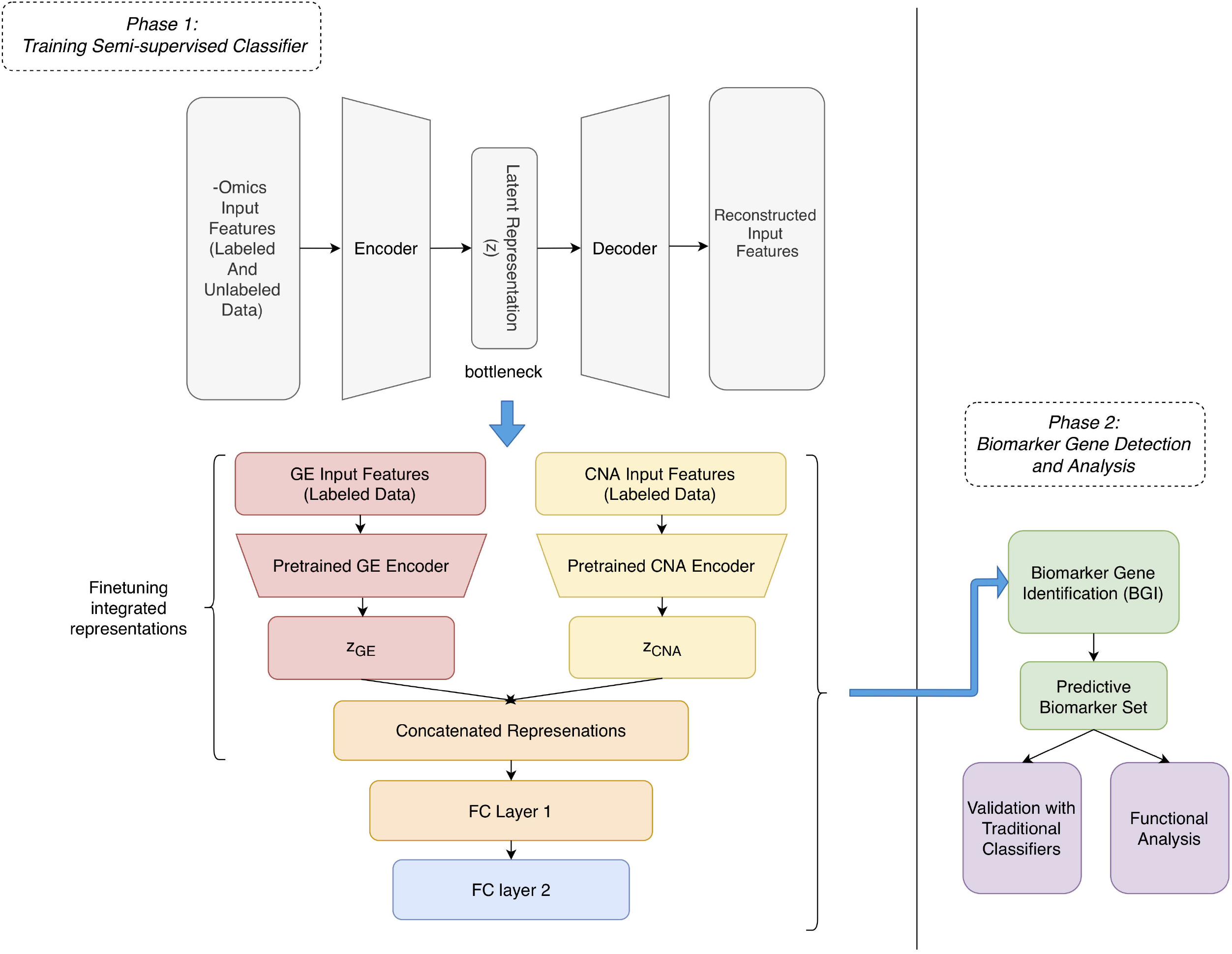
Framework of DeepSSC. In phase 1, the two preprocessed profiles (GE and CNA) from TCGA were fed into their own denoising autoencoders to generate two independent representations, z_GE_ and z_CNA_. Then, z_GE_ and z_CNA_ were concatenated into a single input layer attached with two FC layers to create a neural network classifier. Finally, this neural network classifier became the input of our BGI procedure in phase 2 to detect candidate biomarkers. To demonstrate the efficacy of DeepSSC, those biomarkers should help classify cancer patients into different subtypes well by only using simple machine learning models. Besides, those biomarkers also were further investigated in terms of their biological functions. Note that labeled and unlabeled data were defined clearly at the below Methods section. Abbreviation: GE, mRNA expression; CNA, copy-number alterations; FC, fully-connected layer.

## RESULT

### Semi-supervised classification performance was superior to supervised one

To examine the necessity of with and without additionally the unlabeled samples during the model training process, we performed two experiments using our model on two sub-datasets extracted from the TCGA BRCA dataset. The first sub-dataset consisted of subjects that had their own PAM50 BRCA subtyping scheme in both profiles, GE and CNA, called labeled samples. The second sub-dataset included the first one plus all subjects having their own PAM50 BRCA subtype in at least one type of omics profile or unassigned to any subtype in both GE and CNA data, called unlabeled samples. Consequently, classification performance of the latter case (Figure 2b) was substantially better over that of the former case (Figure 2a) in the three evaluation metrics (i.e., overall accuracy, macro-averaged F1 score, and weighted average F1 score). The same processes and results were done and shown on the COADREAD dataset (Figures 2c,d). These results proved that we could reach a more accurate prediction when using the unlabeled data additionally instead of excluding them.

**Figure 2.**
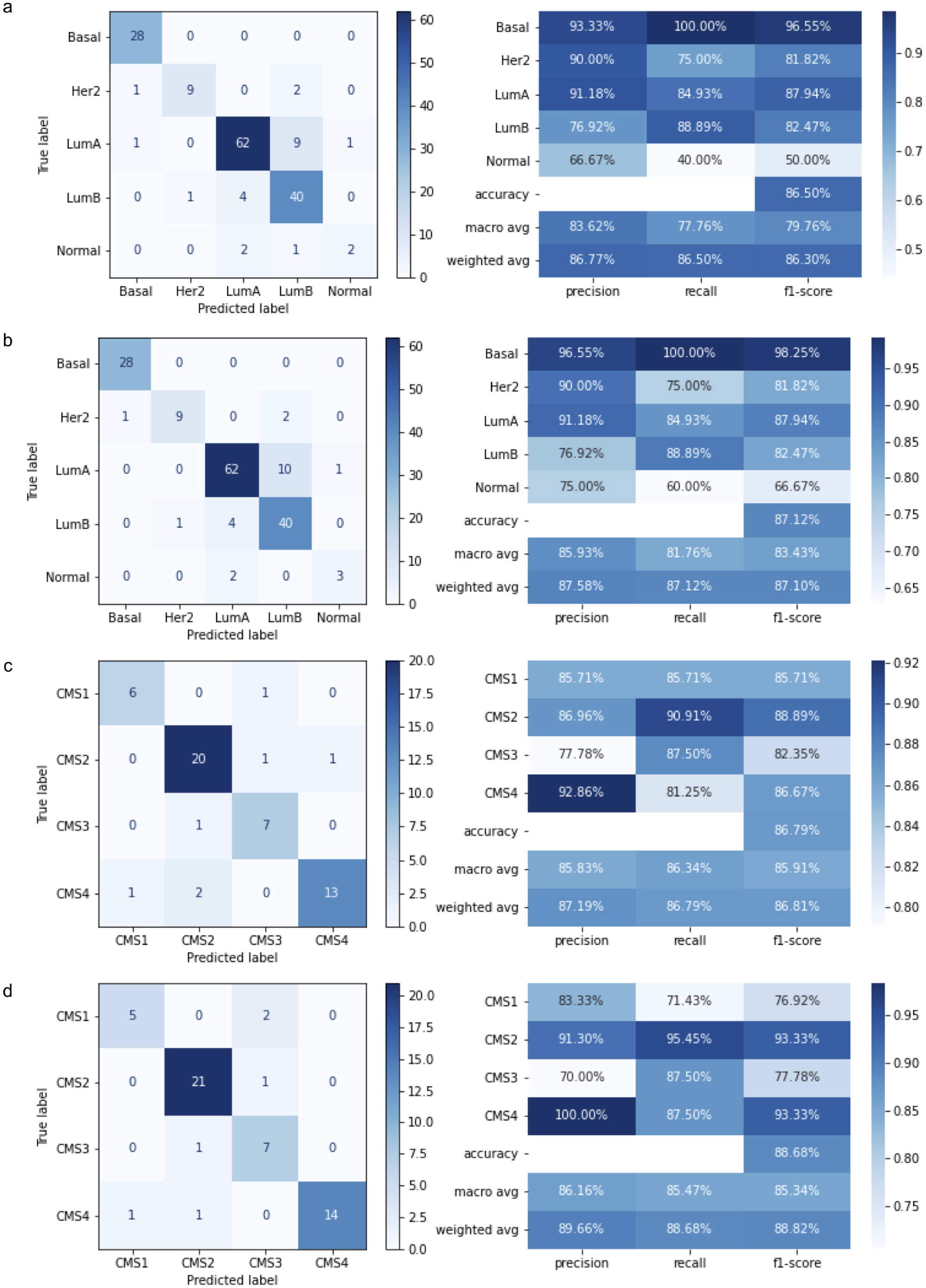
Classification performance of the model using the two different BRCA sub-datasets and two different COADREAD sub-datasets. The model was run individually in the two separate circumstances, supervised learning (a and c) and semi-supervised learning (b and d). **a, c** Model training results for sub-dataset 1 having the only labeled samples in BRCA and COADREAD, respectively. **b, d** Model training results for sub-dataset 2 having both labeled and unlabeled samples in BRCA and COADREAD, respectively. The model architecture and all settings in the model training process were the same for each cancer.

Due to the imbalance of label distribution in BRCA, we tried several weighting strategies to assign weight to labels (as described in the Methods section). As a result, we realized that the forth strategy, i.e., we manually set weight to subtypes, was the best. Table 1 shows our model gained 87.12% accuracy, 83.43% macro F1-score on average, and 87.10% weighted F1-score on average over the five PAM50 classes for integration of GE and CNA. Detailed results for the first three weighting strategies on the BRCA dataset were reported in Supplementary Materials. For the COAREAD dataset, we applied the balance weighting strategy to assign weight to its labels and then obtained results of 88.68% accuracy, 85.34% macro F1-score on average, and 88.82% weighted F1-score on average over the four CMS classes for integration of GE and CNA. As TCGA lung cancer dataset did not suffer from the class imbalance problem, we did not apply the weighting strategy to this dataset. As a result, we received the accuracy of 0.9604, F1 score of 0.9596, and the AUC score of 0.9924 for integration of GE and CNA on the lung cancer dataset (Table 1).

**Table 1.** Classification performance with different omics data types on the three datasets. Abbreviation: ACC, accuracy; macro-averaged F1-score; weighted F1, weighted average F1-score; AUC, Area Under the Receiver Operating Characteristic Curve.

|  | TCGA BRCA |  |  | TCGA lung cancer |  |  | TCGA COADREAD |  |  |
| --- | --- | --- | --- | --- | --- | --- | --- | --- | --- |
|  | ACC | macro F1 | weighted F1 | ACC | F1 | AUC | ACC | Macro F1 | Weighted F1 |
| GE+CNA | **87.12%** | **83.43%** | **87.10%** | **96.04%** | **95.96%** | 99.24% | **88.68%** | 85.34% | **88.82%** |
| GE | 86.50% | 82.83% | 86.28% | 93.56% | 93.33% | **99.26%** | 86.79% | **85.98%** | 86.83% |
| CNA | 61.96% | 41.66% | 60.36% | 89.11% | 88.54% | 97.04% | 37.74% | 34.72% | 35.85% |

### Classification performance of DeepSSC under different -omics data types

To demonstrate the effectiveness of our model with different -omics data types, we compared DeepSSC’s classification performance with different choices of the input -omics data on the three BRCA, COADREAD and lung cancer datasets. The original model was modified into two separate models: one using only GE and the other using only CNA. With the model using only GE data, we removed the encoder of CNA, then the classifier was re-tuned (i.e., changing the number of nodes in the hidden layers and weight decay rate in the last layers) on the validation set, and next re-evaluated on the test set. The same process was applied to the classification model using single CNA data. The results were shown in Table 1. We observed that the classification performance was improved by integrating the two -omics data types in term of the accuracy and weighted F1 score on the COADREAD dataset as well as the accuracy and F1 score on the lung cancer dataset. In contrast, the macro-averaged F1 score was slightly better for the GE-used model on the BRCA dataset and the AUC score for the GE-used model on the lung cancer dataset. For the BRCA dataset, the classification performance was improved across all the three metrics by the integration strategy. Also, for the single -omics approach, the GE-used model was better in classification capability compared with the CNA-used model across all the evaluation metrics on both datasets.

### DeepSSC discovered subtype-specific biomarker genes able to help classify cancer subtypes using classical classifiers

Identification of biomarker genes was crucial for understanding the biological, molecular, and cellular mechanisms. For each subtype, we selected the top-100 important genes, rendering a set of 428 unique ones for BRCA, 359 unique ones for COADREAD, and 127 unique ones for lung cancer (some overlap among the subtypes was allowed). A full list of those potential biomarkers with the corresponding rankings was provided in Supplementary TableS1. To demonstrate the effectiveness of a range of top-five through top-100 important genes extracted from each cancer subtype in differentiating cancer subtypes, we used three traditional classifiers, including Logistic Regression (LR), Support Vector Machine (SVM), and Random Forest (RF). In addition, we further evaluated the effectiveness of these genes under different data types (GE only, CNA only, and integration of GE and CNA) using the LR model only (the most interpretable and simplest classifier among the three selected classifiers). Here the three cancer types actually represented the two common classification problems: multi-class classification (BRCA with the five PAM50 subtypes and COADREAD with the four CMS subtypes) and binary classification (lung cancer with the two histologic subtypes). Moreover, for the case study of BRCA, we also performed on independent dataset, METABRIC BRCA.

#### Classification performance of identified biomarkers on BRCA

We firstly implemented DeepSSC on the two BRCA datasets, TCGA BRCA and METABRIC BRCA. As seen in Figure 3a and Supplementary TableS2, on the TCGA BRCA dataset, each traditional classification method using different sets of top genes yielded high accuracy (> 81.00%), in which the accuracy was proportional to the increased number of top genes, i.e., up to the top-10 genes for SVM, the top-20 genes for LR, and the top-30 genes for RF. At best, the accuracy could reach around 92.00% for all of the three classifiers SVM, RF, and LR at the top-10, −30, and −60 genes, respectively. Clearly, irrespective of the simple classification model we chose, our identified biomarkers had the potential to distinguish the five BRCA subtypes well. Besides, as shown in Figure 3b and Supplementary TableS3, we could see that when using only CNA data, the accuracy and macro-average F1 score across all top genes were substantially lower than when using only GE data or integration of the two. Also, as compared to the classification performance of the models using the integrated data, that of the models using only GE data was not extremely inferior at top gene sets.

**Figure 3.**
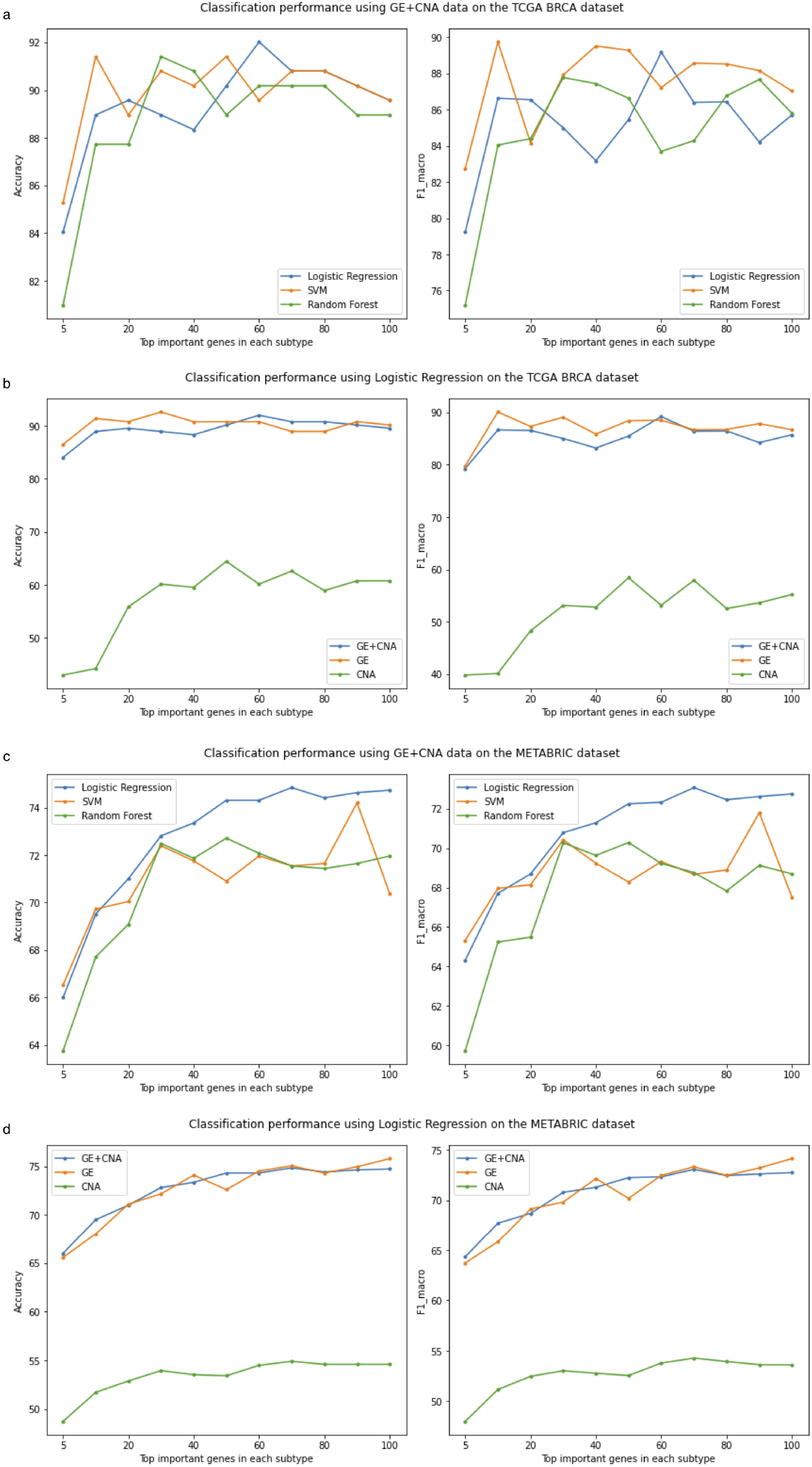
Classification performance comparisons on TCGA and METABRIC BRCA. (a and c) Classification performances using the three classifiers on TCGA (a) and METABRIC (c). We pulled a range of top-5 through top-100 important genes from each intrinsic subtype. The gene sets were then used as the input features for the different classifiers to measure classification performance. (b and d) Classification performances using the Logistic Regression algorithm on different omics data types (integration of GE+CNA, GE, and CNA) from TCGA (b) and METABRIC (d). We pulled a range of top-5 through top-100 important genes from each intrinsic subtype. Each model was run using repeated k-fold cross-validation with k of five and the number of repetitions n of 10. The model with the highest macro-averaged F1 over all folds (k x n) was evaluated on the TCGA test set as well as on the METABRIC validation set.

Moreover, we also validated our candidate gene sets on the independent dataset METABRIC with the same three classifiers as mentioned above. In Figure 3c and Supplementary TableS4, we could see that the best accuracy of each classifier was fairly high (> 72.00%), and it could reach the accuracy of 74.84% with only a simple classifier as the LR algorithm. Also, the accuracy of the three classifiers was proportional to the increase in the number of top genes, which was evident most in the LR classifier. These demonstrated that the 428 predictive biomarker genes were still valuable in terms of the BRCA subtype classification independent of patient cohort and data platforms. Futhermore, Figure 3d and Supplementary TableS5 show the performance pertaining to both accuracy and macroaverage F1 score across the list of various top gene sets using the different -omics types.

#### Classification performance of identified biomarkers on COADREAD

We secondly implemented DeepSSC on the TCGA COADREAD dataset, another example for the multi-class classification problem. As shown in 4a and Supplementary TableS6, the accuracy of all of the three classifiers was above 86% for all top genes. The simplest classifier LG still issued feasible results using our identified biomarkers and could achieve the highest accuracy of about 98% in the top-50 genes (same results as SVM). Futhermore, Figure 4b and Supplementary TableS7 show the performance pertaining to both accuracy and macro-average F1 score across the list of various top gene sets using the different -omics types. These indicated that DeepSSC could work well with the multi-class classification problems.

**Figure 4.**
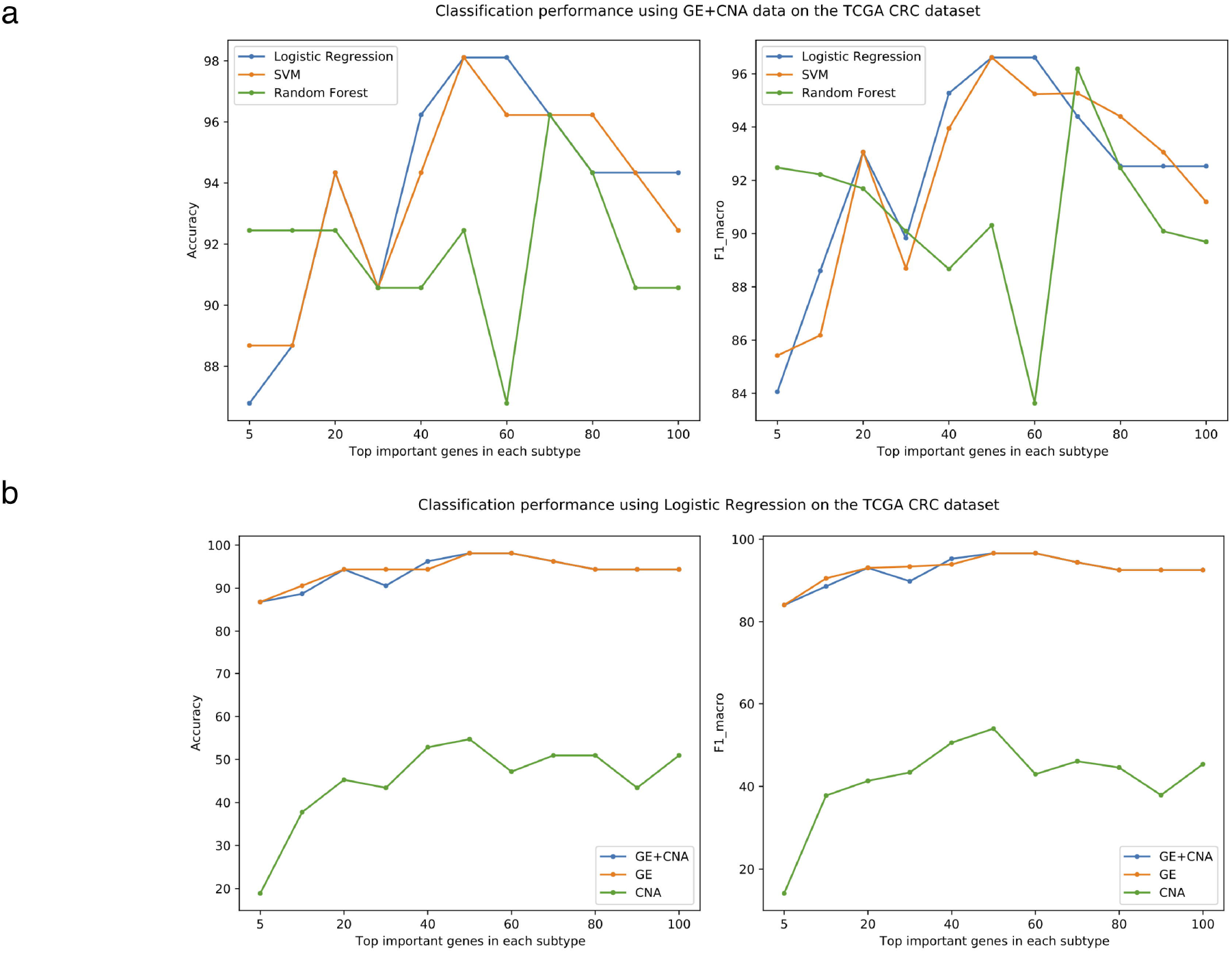
Classification performance comparisons on TCGA COADREAD. (a) Classification performance using the three classifiers. We pulled a range of top-5 through top-100 important genes from each CMS subtype. The gene sets were then used as the input features for different classifiers to measure classification performance. (b) Classification performance using the Logistic Regression algorithm on different omics data types (integration of GE+CNA, GE, and CNA). We pulled a range of top-5 through top-100 important genes from each CMS subtype. Each model was run using repeated k-fold cross-validation with k of five and the number of repetitions n of 10. The model with the highest accuracy over all folds (k x n) was evaluated on the test set.

#### Classification performance of identified biomarkers on lung cancer

Lastly, we also validated the ability of DeepSSC to differentiate the lung cancer subtypes, a simpler classification problem (binary classification). The accuracy and F1 score were quite similar, so we only showed the accuracy and AUC scores here (all detailed information about the accuracy, F1 scores, and AUC scores were described in the Supplementary TableS8). As we could see in Figure 5a and Supplementary TableS8, with only the top-five genes per subtype, including six unique genes in total, all of the three traditional classification methods could reach high performance in both accuracy and AUC metrics (>94.5% accuracy and >98.4% AUC). Performance of the three classifiers did not change too much when increasing the number of top genes per subtype. The gap between the lowest and highest performance of each classifier only fluctuated from 1% to 2% accuracy and <1% AUC score. More noticeably, the model’s performance was high at the top-five gene set and minimally changed when going through the following top gene sets. Besides, as shown in Figure 5b and Supplementary TableS9, we once again found that the classification performance of the models using integration of GE and CNA was similar to that of the models using only GE data, and higher than that of the models using only CNA data. These pointed out that DeepSSC could be used effectively for cancer subtype classification if even only using the tiny number of top genes per subtype. In addition, DeepSSC could also possibly work well for both binary and multi-label classification problems.

**Figure 5.**
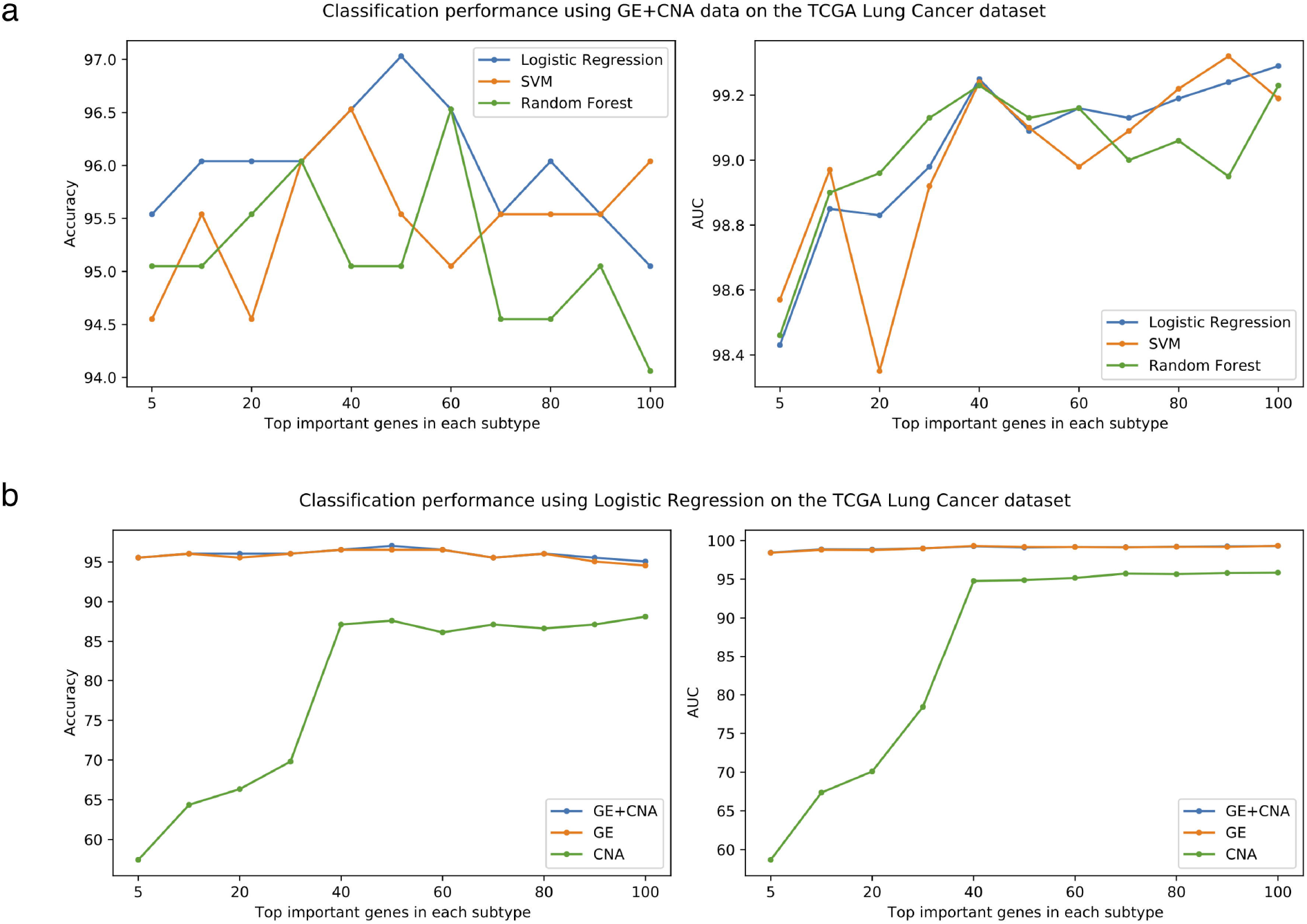
Classification performance comparisons on TCGA lung cancer. (a) Classification performance using the three classifiers. We pulled a range of top-5 through top-100 important genes from each histologic subtype. The gene sets were then used as the input features for different classifiers to measure classification performance. (b) Classification performance using the Logistic Regression algorithm on different omics data types (integration of GE+CNA, GE, and CNA). We pulled a range of top-5 through top-100 important genes from each histologic subtype. Each model was run using repeated k-fold cross-validation with k of five and the number of repetitions n of 10. The model with the highest accuracy over all folds (k x n) was evaluated on the test set.

### DeepSSC outperformed other state-of-the-art methods

We saw the effectiveness of semi-supervised learning in multi-omics classification so far (Figure 2). However, we wondered whether the structure of DeepSSC was optimal or not. Accordingly, DeepSSC concatenated latent-space representations of -omics data after they were put into DAE. To examine the necessity of post-concatenation strategy, DeepSSC’s three additional variations were compared: (i) pre-concat-AE: directly concatenating the input -omics data features before being put into AE, (ii) pre-concat-DAE: directly concatenating the input -omics data features before being put into DAE, and (iii) pre-concat-VAE: directly concatenating the input -omics data features before being put into variational AE (VAE). The three variations of DeepSSC were implemented with our semi-supervised strategy, and with the same number of layers and the same dimensions of hidden layers as the DAE part in DeepSSC. In practice, these variations were commonly encountered in previous studies, such as [19, 22]. Furthermore, we also sought to compare the classification performance of DeepSSC with a recently proposed state-of-the-art supervised multi-omics integrative analysis method named MOGONET. MOGONET [17] was a deep learning framework using the graph convolutional networks to explore omics-specific learning and then using View Correlation Discovery Network to explore the cross-omics correlations at the label space. Furthermore, MOGONET used an ablation approach (i.e., set each feature’s value to 0, then, the feature importance would be measured by the performance decrease after the feature was masked) to identify important biomarkers. Thus, this model essentially required multi-stages feature preselection. More specifically, we compared DeepSSC with its variations and MOGONET in light of the accuracy, weighted F1 score, and macro F1 score for the BRCA and COADREAD datasets, and the accuracy, F1 score, and AUC for the lung cancer dataset. At phase 2, we only inputted the top-10 important genes within each subtype, corresponding 46, 38 and 14 genes in total for BRCA, COADREAD and lung cancer, into the three classifiers (SVM, LR, and RF).

As shown in Table 2, we observed that DeepSSC showed a substantially better classification performance over its variations. These results indicated that the use of post-concatenation strategy in DeepSSC was more effective than that of the pre-concatenation strategy under the semi-supervised learning condition in biomedical classification problems.

**Table 2.**
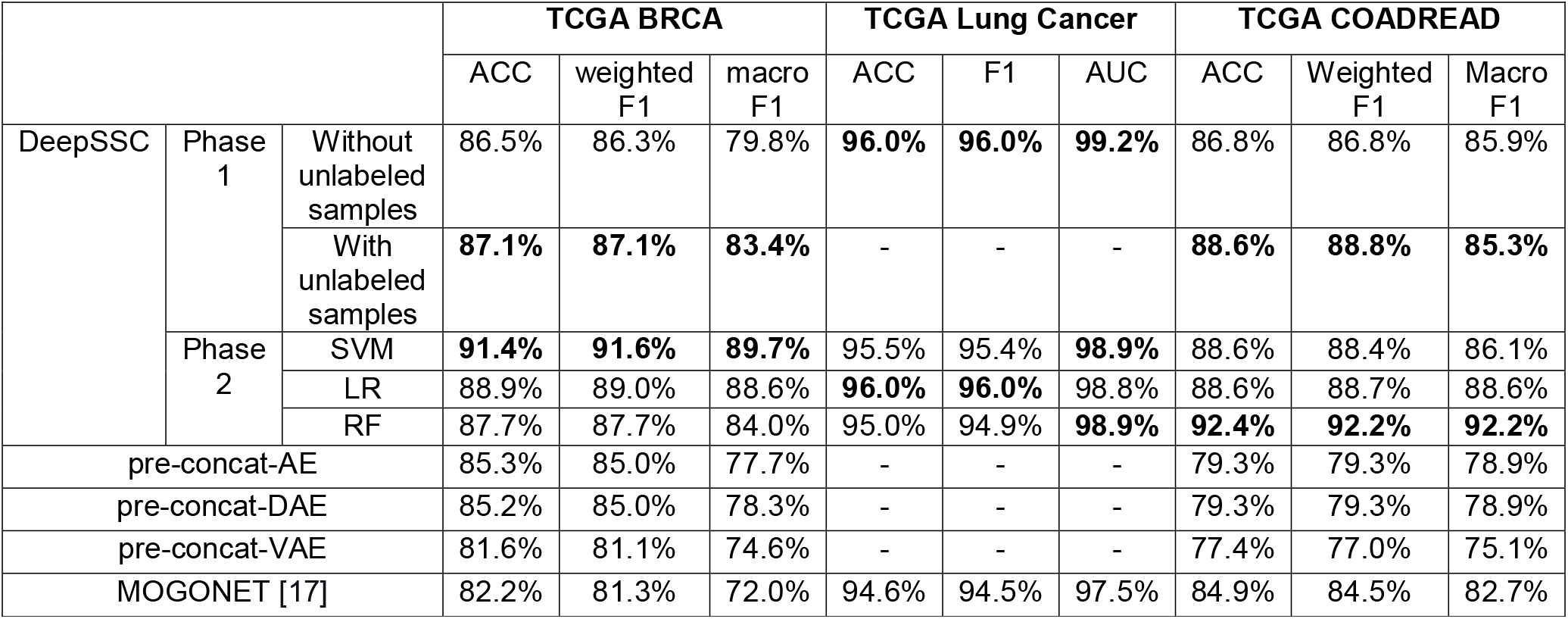
Comparison of DeepSSC with the state-of-the-art techniques for BRCA subtype classification, COADREAD subtype classification, and lung cancer type classification. We only inputted the top-10 important genes per subtype into the three classifiers (SVM, LR, and RF) at phase 2. We rounded 0.5 to the next nearest whole number.

Moreover, DeepSSC also outperformed MOGONET in all classification tasks (Table 2). Indeed, BRCA-specific DeepSSC, COADREAD-specific DeepSSC, and lung-cancer-specific DeepSSC consistently achieved the best results in different metrics at phase 1. For the second phase, it was of great interest that all of the simple machine learning models of our choice exhibited better performance over MOGONET on all of the three cancer types. Clearly, this demonstrated that DeepSSC had the superiority of semi-supervised multi-omics data classification capability even if solely using the simple machine learning techniques alongside the minimum number of input features.

### DeepSSC discovered BRCA-associated biomarkers

Table 3 shows a set of important biomarkers considered to be genuine in BRCA, in which several well-known genes ranked high in the list such as *GRB7, ERBB2, PIK3R1,* etc. To be consistent with the number of PAM50 genes indicated previously by Parker *et al*.[23], we also picked out the 50 top-ranked biomarkers (top-10 genes per intrinsic subtype; bold red gene names in Supplementary TableS1a) to undergo biologically functional analyses as described in the Methods section. As a result, we gained a set of 46 genomic biomarkers in total, in which *C10orf26, WDR67,* and *C9orf100* were overlapped one or more times among the subtypes. Figure 6a illustrates the heatmap showing differences in expression levels of these biomarkers among the five molecular subtypes. Three genes, including *ERBB2, MIA,* and *GRB7*, were included in the BRCA multi-gene signature.

**Table 3.** Comparisons between predictive biomarker genes and four well-known sources of BRCA-associated genes.

| Top-100 genes in each subtype | Cancer Gene Census [34] | Nik Zainal *et al.* paper [35] | Pereira *et al.* paper [21] | PAM50 genes [23] |
| --- | --- | --- | --- | --- |
| Basal | FOXA1, GATA3 | FOXA1, GATA3 | GATA3 | FOXA1, MAPT, MIA |
| Her2 | ERBB2 | ERBB2, FGFR2 | ERBB2 | BCL2, ERBB2, GRB7 |
| LumA | - | - | - | ANLN, BCL2, EXO1, MAPT, MELK |
| LumB | - | - | - | ANLN, CDC6, KRT17, KRT5, MIA |
| Normal | - | PIK3R1 | - | - |

**Figure 6.**
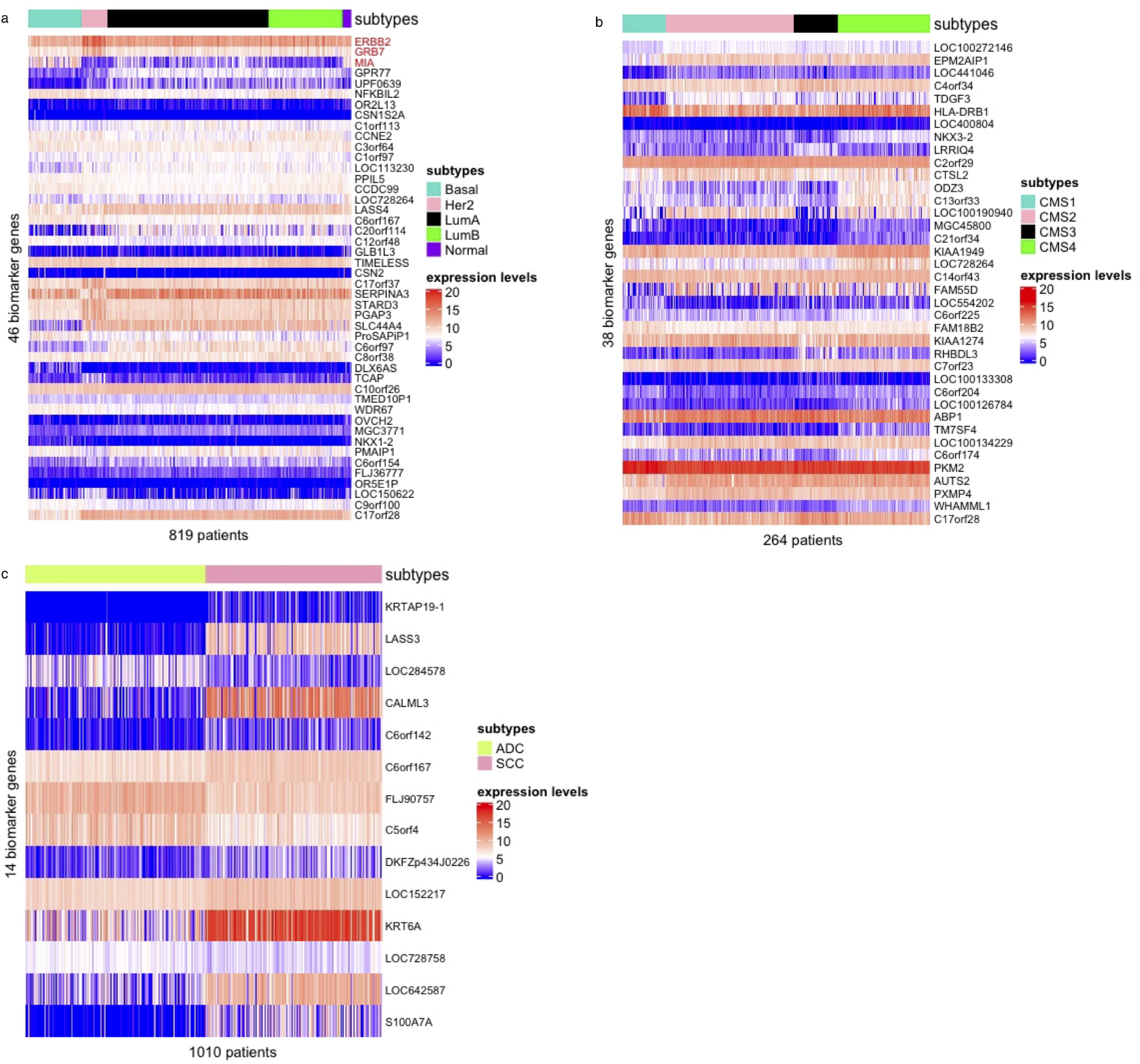
Heatmaps of expression leves of selected biomarker genes among subtypes. (a) Heatmap showed differences in expression levels of BRCA biomarkers among the five PAM50 subtypes. Red gene names indicated that they were included in the BRCA PAM50 signature. (b) Heatmap showed differences in expression levels of biomarkers among the four CMS subtypes. (c) Heatmap showed differences in expression levels of biomarkers between the two lung cancer subtypes. Abbreviation: node, the lymph node and size, the size of the tumor; ADC, lung adenocarcinoma; SCC, lung squamous cell carcinoma.

Firstly, we noticed an interesting result that two highly-ranked genes, *ERBB2* and *GRB7,* assigned computationally to the HER2 subtype by DeepSSC, were associated highly with BRCA as well as this molecular subtype. The gene *GRB7* was indeed located in close proximity to the *ERBB2* and they overexpressed with each other in BRCA tumors [15, 23]. Next, we did enrichment analysis on the 46 predicted genes to discover their specific biological meaning. As a results, the enriched pathways within these signatures were also discovered (Supplementary Table S10). Some cancer-associated pathways were indicated, including p53 signaling pathway (hsa04115; adjusted P-value□=□0.03), prostate cancer (hsa05215; adjusted P-value□=□0.04), etc. These results further confirmed that the 46 biomarker genes predicted by DeepSSC had key functions in cancer in general and BRCA in particular.

### DeepSSC discovered COADREAD-associated biomarkers

Similar to the way to select biomarkers in BRCA, we also pulled the 40 top-ranked COADREAD-associated biomarkers from Supplementary TableS1b (top-10 genes per CMS subtype; bold red gene names). As a result, we gained a set of 38 genomic biomarkers in total, in which the two genes *ODZ3,* and *WHAMML1* were overlapped among the subtypes. Figure 6b illustrates the heatmap showing differences in expression levels of these biomarkers among the four CMS subtypes. As a result, we found that several statistically significant terms showed demonstrably closed relationships between COADREAD versus type I diabetes mellitus (hsa04940, adjusted P-value = 0.04, Supplementary Table S11) [24], versus autoimmune thyroid disease (hsa05320, adjusted P-value = 0.04, Supplementary Table S11) [25, 26], and versus Inflammatory bowel disease (hsa05321, adjusted P-value = 0.04, Supplementary Table S11) [27].

### DeepSSC discovered lung-cancer-associated biomarkers

We also pulled the 20 top-ranked lung-cancer-associated biomarkers from Supplementary TableS1c (top-10 genes per subtype; bold red gene names). As a result, we gained a set of 14 genomic biomarkers in total, in which the six genes *LASS3, FLJ90757, LOC284578, CALML3, KRTAP19-1*, and *LOC152217* were overlapped among the subtypes. Figure 6c illustrates the heatmap showing differences in expression levels of these biomarkers between the two histologic subtypes. Consequently, we found that the statistically significant term was keratinocyte differentiation (GO:0030216, adjusted P-valuel□=□10.008, Supplementary Table S12) associated with lung cancer [28]. In addition, cytolysis (GO:0019835; adjusted P-valuel□=□10.01, Supplementary Table S12), playing a vital role in the immune system, also was reported to be useful for building new approaches for the treatment of lung cancer [29].

## DISCUSSION

In the past, the collection and annotation of data have been highly expensive plus knowledge of disease subtyping has been limited, resulting in lack of the labeled biomedical data. Therefore, most studies using or proposing multi-omics integration methods or tools will concentrate only on unsupervised learning algorithms without additional phenotypic information, and then attempt to gain useful biological information from the identified sample clusters. Nevertheless, due to the unprecedented proliferation of omics technologies and the establishment of cancer genomic databases such as TCGA and METABRIC, labeled -omics datasets with detailed annotations have been publicly available. Consequently, it has enabled cancer researchers to leverage these labeled -omics data to better predict essential phenotypes or clinical features on new samples. However, the number of labeled samples alone is still quite small for learning capacity of modern deep learning models. With these concerns in mind, we propose a biphasic Deep Semi-supervised multi-omics integration framework for Subtype Classification and biomarker discovery, DeepSSC, based on deep multi-aspect learning where we regard each -omics data to be an aspect of the samples. We utilize DAE for omics-specific representation learning, while a multi-layer perceptron for multi-omics representation integration and subtype classification. To this end, we envision DeepSSC as a novel deep learning framework for classification tasks that takes both superior performance and good interpretability.

In comparison with the classification performance of DeepSSC with and without unlabeled samples (Figure 2), we demonstrate that semi-supervised learning is needed for effective multi-omics data classification. Moreover, by the comparison of the classification capability between DeepSSC and its three variations, we also demonstrate that DeepSSC effectively classifies multi-omics data by concatenating the omics-specific representations generated by DAE at the latent space, called the post-concatenation strategy in this work. A possible explanation for these outperformances is that the pre-concatenation approach is more likely to be biased toward certain -omics data types rather than the post-concatenation approach. Another explanation for these using the pre-concatenation approach would be unable to leverage the power of semi-supervised learning because not including several patients whose information only belonging to the one input data but not to the other input data, resulting in loss of potentially useful information. For example, only one of the input -omics data having no unlabeled samples, then all of unlabeled samples cannot be included in the model pretraining. Besides, the BGI procedure is used to identify subtype-specific biomarker genes in various cancer types. They are useful to help to classify cancer subtypes highly accurately even if only using classical machine learning techniques with a small quantity. We also further indicate that the subtype-specific biomarkers have a good biological correlation with each certain cancer type. Additionally, we, in this study, apply DeepSSC to the three case studies of BRCA, COADREAD, and lung cancer as well as make use of the GE and CNA data for multi-omics classification task; however, it can be potentially applicable to distinct or more types of data of any cancer type (e.g., kidney cancer, brain lower grade glioma, etc.) or any complex disease (e.g., Alzheimer’s disease).

Figure 2 shows a limitation of DeepSSC that it has trouble discriminating BRCA LumA and LumB tumors separately. One potential explanation comes from the label imbalance in the data. Moreover, the PAM50 classification system is expected to be more robust than other conventional ones based solely on a few biomarkers, the inconsistency of discrimination between LumA and LumB by the various predictors, suggesting that these label classes may not represent distinct coherent sample groups [30]. Because of this problem, we [3] and Dvir Netanely *et al.* [9] have attempted to recluster patients with these two BRCA subtypes. Looking back through history of BRCA classification, luminal tumors have also experienced the re-classification many times. From that, we suggest that the future works should find out a better solution for this problem.

To sum up, DeepSSC successfully and effectively classifies cancer subtypes, detects genuine biomarkers as well as hints at novel candidates from integrated -omics data types that show strong associations with BRCA, COADREAD and lung cancer. Through this study, we wish to send a message that we should take adoting the semisupervised techniques into account in -omics-based classification problems.

## METHODS

### Data collection and preprocessing

#### Data collection

The Cancer Genome Atlas (TCGA) used in the study were extracted from the UCSC Cancer Genome Browser Xena [31]. METABRIC BRCA dataset [21] was extracted from cBioPortal [32, 33] and split into discovery and validation sets according to European Genome-Phenome Archive (EGA) under the accession number EGAS00000000083. The CNA (measured on the Affymetrix SNP 6.0 platform) and GE (generated using Illumina Human v3 microarray) data were in the same format as the TCGA data.

#### Data preprocessing

All of the datasets obtained from TCGA underwent a process for which we eliminated invalid samples if they existed: solid tissue normal and metastasis samples. Here the labeled data were samples assigned to a specific disease subtype in both the GE and CNA data. In contrast, the unlabeled data were samples assigned to a specific disease subtype in only either of the GE or CNA data or unassigned to any specific disease subtypes in both GE and CNA data (Supplementary Figure S1). Instead of removing the unlabeled data, we used these data to take full advantage of information during training autoencoders for the TCGA BRCA and TCGA COADREAD datasets. Besides metastatic and solid tissue normal samples, some BRCA patients which were male or included in the blacklist (http://gdac.broadinstitute.org/runs/stddata_2014_02_15/samples_report/Blacklisted_Samples.html) and/or redaction list (http://gdac.broadinstitute.org/runs/stddata_2014_06_14/samples_report/BRCA_Redactions.html) did not guarantee the quality for our research purpose, and so were removed from the original dataset.

As a result, in the TCGA BRCA dataset, we received 819 labeled samples for both -omics data, and 259 and 245 unlabeled samples for the GE and CNA data, respectively. Out of 819 labeled samples, 409 for the LumA subtype, 187 for the LumB subtype, 134 for the Basal-like subtype, 67 for the HER2 subtype, and the remaining 22 samples for Normal-like subtype. Similarly, in the TCGA COADREAD dataset, we obtained 264 labeled samples for both -omics data, and 118 and 352 unlabeled samples corresponding to the GE and CNA data. The four TCGA COADREAD CMS subtypes were received from [34]. We received 1,010 labeled samples for both -omics data for the TCGA lung cancer dataset. We did not use the unlabeled data because of its small sample size. The summary of each subtype corresponding to the TCGA/METABRIC BRCA, TCGA COADREAD, and TCGA lung cancer were shown in Table 4. Finally, the labeled data in all TCGA datasets were further randomly divided into three sets, including training, validation, and test sets, in the proportion of 64:16:20, respectively. It should be noted that we retrieved sample statistics (i.e., mean and standard deviation) of the training set in case study of TCGA lung cancer or of the data, including the labeled data of the training set and the unlabeled data in case studies of BRCA and COADREAD, and applied them to Z-normalization to normalize all of the three sets before training any neural networks.

**Table 4.**
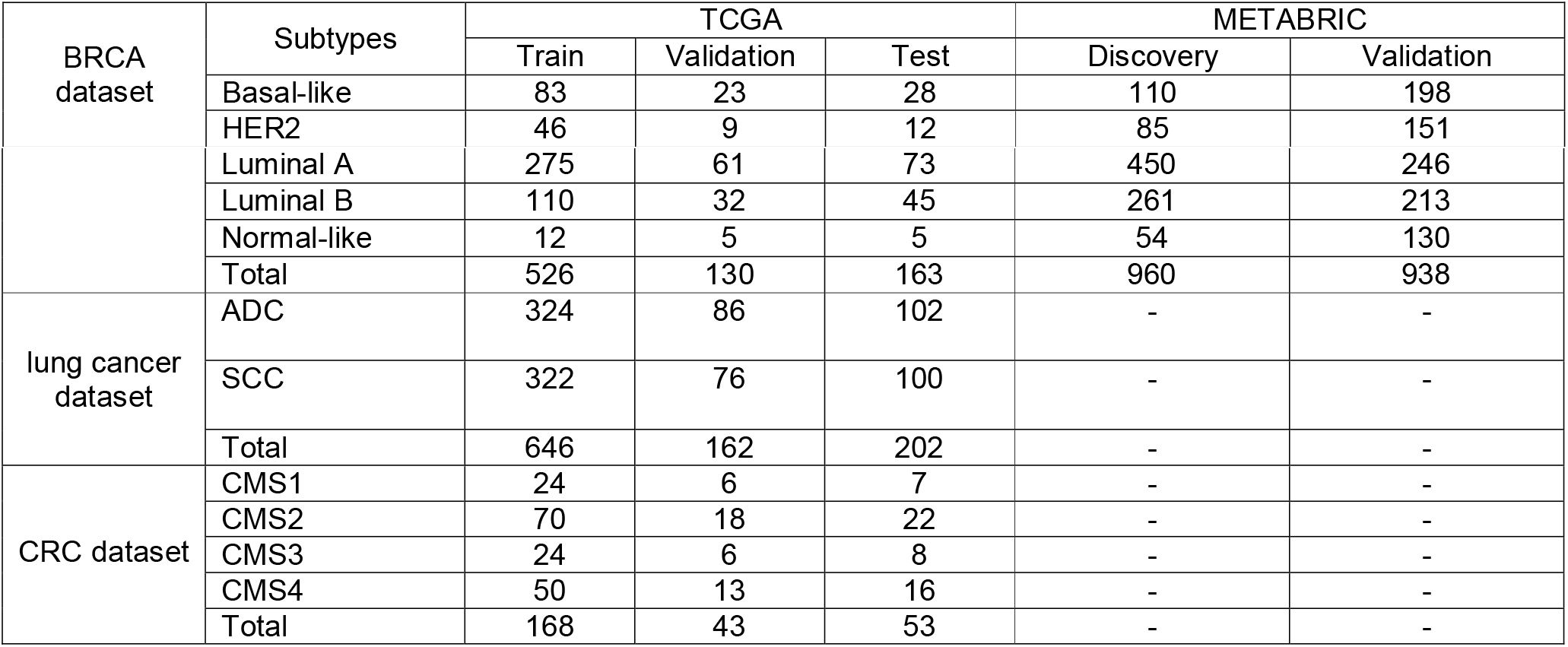
Data summary of TGCA/METABRIC BRCA, TCGA COADREAD, and TCGA lung cancer. Abbreviation: BRCA, breast cancer; ADC, lung adenocarcinoma; SCC, lung squamous cell carcinoma.

We applied the same pre-processing procedure to the discovery and validation sets of the METABRIC dataset. Note that for two or more genes sharing the same name in the data (i.e., the same “Hugo Symbol Entrez” and “Gene_ID” fields), their average values were used to merge them. Therefore, we gained 17,300 genes for GE data and 22,544 genes for CNA data. At last, we only kept the labeled samples with biomarkers were discovered by the BGI procedure in the TCGA dataset in both CNA and GE data to train BRCA subtypes classifiers. The three classical machine learning techniques were used on the discovery set and then classification performance were evaluated on the validation set.

#### Training semi-supervised neural network for classification tasks

Due to the different nature of the two data types, we built two independent DAE for the two TCGA datasets (Figure 1). Specifically, each DAE was trained with each -omics data including its labeled and unlabeled samples. DAE for GE data consisted of the five FC layers with exponential linear unit (ELU) (40) activation functions. The ELU activation function was calculated as follows:

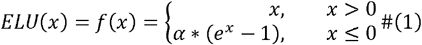

The first three layers were called the encoder including the input layer and two hidden layers with dimensions of 20,530, 4,096, and 2,048, respectively. The following two layers, called the decoder, had the number of hidden nodes was 4,096 and 20,530, respectively. likewise, DAE for CNA data also consisted of the five FC layers with dimensions of 24,776, 4,096, 1,024, 4,096, and 24,776, respectively. The model’s hyper-parameters (number of layers *L,* number of hidden units *H,* and corruption rate *p)* were chosen so that the Mean Square Error (MSE) on the validation set was minimal. We found that the optimal drop-out rate *p* was 0.5 on the input vector in this case. To deal with the internal covariant shift problem, a batch normalization layer was applied before the activation function at the first hidden layer of each encoder part.

The encoders from the two DAEs described above were used as pretrained models for the subtype classification network. The outputs of the two encoders were then concatenated into a single 3,072-dimensional feature layer. This layer was attached with a hidden layer containing 1,024 neurons. The final neural network classifier consisted of the two encoders and the two FC layers with ELU as their activation functions. The last FC layer was called the output layer, including the class scores representing each cancer subtype. The pretrained weights of encoders were then fine-tuned with smaller learning rate than those of two last layers to bring the representations of the two omics data types to the same representation space being suitable for the classification task. The model’s hyper-parameters (*H* and *p*) were chosen so that the accuracy on the validation set was maximal. We used the early-stopping technique when training the classifier to get rid of the overfitting problem. To deal with the class imbalance problem, we implemented a weighted cross-entropy loss as described in detail in the next section.

#### Weighted-Cross entropy for imbalanced dataset

TCGA BRCA data was heavily unbalanced because the number of samples varied in each subtype; e.g, normal-like samples were too small compared to other labels after our rigorous pre-processing step. To deal with this problem, we tried several weighting strategies for multi-class crossentropy loss. The most common strategy was a balanced heuristic inspired by [35]. The weight was calculated as follows:

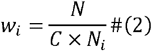

where *N* total number of samples, *C* number of classes, *w_i_* weight for class *i*.

To prevent a large difference between the weight of the minority class like the normal-like label and the weight of the majority class like the LumA label, from negatively affecting model performance, we converted *w_i_* in **Eq. 1** to a logarithmic scale as follows:

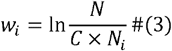

Although we saw the efficacy of the two strategies above, we attempted to continue to improve the classification performance more. Indeed, we manually adjusted the weight of each subtype based on the recall and f1-score in classification result of each class on the validation set, at which point these weights were treated as a hyperparameter of the model. The weights of Basal-like and HER2 classes were unchangingly set to one due to their ease of classification compared to other classes, possibly the number of samples in these two labels was large most, the weights of the remaining subtypes were gradually increased so that the average F1-scores on the validation set was maximal. These weights were then applied to the cross-entropy loss function as follows:

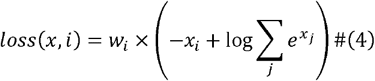

The losses were averaged over observations for each mini-batch as follow:

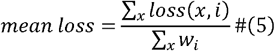

where *x* predicted probability distribution over classes of sample *x, i.* ground truth class of *x.*

#### BGI procedure

Our aim of the BGI procedure was to identify genes that played a significantly important role within each subtype. More specifically, the BGI procedure used the trained neural network classifier as the input of the Captum tool [36] (version 0.4.0), a unified and generic neural networks model interpretability tool, to find biomarker genes. This tool included state-of-the-art algorithms to interpret many types of models and understand which features were important, in which an attribution method called integrated gradients was selected in the study. Given a certain output of the neural network (target neuron), the algorithm computed the gradient of the prediction output of the network to mark the features (i.e., genes) that played a role in arriving at the output of the model. More detailed information of BGI procedure was exhaustedly described in the Supplementary Methods section in Supplementary Materials.

The integrated gradients algorithm, the i^th^ dimension (i.e., i^th^ gene) for an input features x (i.e., GE and CNA) and the baseline x’ was defined as follows:

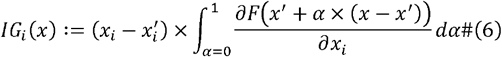

where the integral of the integrated gradients (IG) could be efficiently approximated via a summation. This could be done by simply summing the gradients at points occurring at sufficiently small intervals along the straight line path from the baseline x’ to the input x.

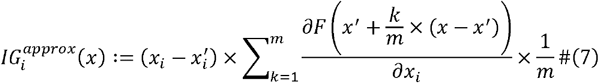

where *m* the number of steps in the Riemman approximation of the integral. Notice that the approximation simply involved computing the gradient in a for loop which should be straightforward and efficient in most deep learning frameworks.

In addition, to verify BRCA-related biomarkers predicted by our BGI, we referred to the four ground-truth sources, including the Cancer Gene Census database (https://cancer.sanger.ac.uk/census) [37], Pereira *et al.* paper [21], Nik Zainal *et al.* paper [38], and the PAM50 genes [23]. Finally, those biomarkers were said to be *bona fide* ones if they appeared in one of the sources above.

#### Functional analysis of predictive biomarkers

To functionally analyze predictive biomarkers, we used clusterProfiler (version 3.16.1) [39] to discover involved biological processes and KEGG pathways. GO terms and KEGG pathways were considered to be significantly enriched if adjusted P-value□≤□0.05 (Benjamini-Hochberg procedure [40], two-sided).

#### Environment and hyperparameter settings

All models were built in Python 3.7.9 on the Kaggle platform with four CPU cores, 16GB Ram for CPU usage, and two CPU cores, 13 GB Ram, 16GB VRAM for NVIDIA Tesla P100 GPU usage. We used the Adam optimizer method [41] with an initial learning rate of 10^-4^ to train the autoencoder and learning rate of 10^-5^ and 10^-6^ for classification part and pretrained part of the fusion model. In both cases, we set a batch size to 32. The functional analyses were performed under R version 4.0.0 on a Mac OS High Sierra 10.13.6 with 8 GB of RAM.

## DATA AVAILABILITY

The raw data from TCGA and METABRIC used in the study are available in the UCSC Cancer Genome Browser Xena (https://xenabrowser.net/) and the cBioportal website: (http://www.cbioportal.org), respectively. Approval by a local ethics committee was not required, and all the data can be immediately downloaded to serve for research purposes. We also made them available by pushing to this (https://www.kaggle.com/dataset/a63f5d7411c02e9aa222ddc53384d342a721b06c2d59cba94e1bb68cf66a535f).

## CODE AVAILABILITY

Python and R codes for reproducing the all results shown in the study (...).

## Author Contributions

Q.-H.N conceived and designed the approach. H.L, V.-M.N, and Q.-H.N drafted the manuscript, which was edited by D.-H.L. H.L and V.-M.N built and ran the models in Python. Q.-H.N performed the analyses in R and interpreted the results. D.-H.L provided critical feedback and key elements during the study period. Q.-H.N and D.-H.L jointly directed and supervised the work. All authors read and approved the final manuscript.

## Correspondence

Correspondence to Quang-Huy Nguyen or Duc-Hau Le.

