## Supplementary Materials for "A biphasic Deep Semi-supervised framework for Suptype Classification and biomarker discovery"

### 1. Supplementary Methods

#### 1.1. Data pre-processing

Instead of removing the unlabeled data, we use all these data to take full advantage of information during training autoencoders in case studies of TCGA BRCA and TCGA COADREAD. As shown in Figure S1, the labeled data are samples assigned to a specific subtype in both the GE data and CNA data. In contrast, the unlabeled data were samples assigned to a specific subtype in either of the GE or CNA, or unassigned to any specific subtype in both the two GE or CNA.

#### 1.2. BGI procudure

In the second phase of DeepSSC, the aim is to identify the genes that have a salient contribution in achieving the classification results. For this purpose, we have proposed the Biomarker Gene Identification (BGI) procedure that leverages the neural network classifier of the first phase for detecting important biomarker genes using relevance propagation methods available in the Captum tool. Specifically, we have selected the algorithm Integrated Gradient. This method is used for reasoning the behavior/outcome of neural network. Given a certain output of the neural networks, this method backpropagates to mark the features (i.e., *candidate_genes*) that play a significant role in arriving at the output of the neural network, thus it is also called a backpropagation method. For a given subtype (say, Basal-like), we select the top-100 important genes that contribute to this subtype. Due to the nature of the binary classification problem, genes that contribute positively to the first subtype will often contribute negatively to the second subtype. Therefore, to this problem, to ensure the target number of top genes, the top genes per subtype will be selected so that genes in the same subtype are unique, while some overlap between the two subtypes was allowed. The sets of selected genes corresponding to the different subtypes were merged into a single set of genes, called *candidateGenes*. The procedure of BGI is described in algorithm 1.

ALGORITHM 1: BGI algorithm

INPUT:	W: Trained neural network model.

C : Subtypes label.

D : All patients have true subtype prediction by W in training and validation set, size MxN, where M denotes number of patients and N denotes number of genes.

k: Threshold k denote number of genes will get for each subtype for further analysis.

OUTPUT: Top k most importance genes each subtypes

METHOD:

for each subtype $p$ in subtypes do:

% get the index of patients whose subtypes are labeled as subtype $i$ and other

% subtypes

$S_{p}\leftarrow\left\{ j|C_{j}=p \right\}$;

$\bar{S}_{p}\leftarrow\left\{ j|C_{j}\neq p \right\}$;

end

% $candidate\_genes_{p}$ denotes top k genes by their contribution scores to subtype $p$

$$candidate\_genes\leftarrow\emptyset$$

for each subtype $p$ in subtypes do:

% ${att\_scores}_{ij}, \bar{att\_scores}_{ij}$ denote attribution scores of gene $j$ of patient $i$ with respect % to subtype $p$ from data set $D_{S_{p}}$ and $D_{\bar{S}_{p}}$.

$att\_scores\leftarrow IntegratedGradients\left( W, D_{S_{p}}, p \right)$;

$\bar{att\_scores}\leftarrow-1\times IntegratedGradients\left( W, D_{\bar{S}_{p}}, p \right)$;

% ${gene\_scores}_{j}$ denotes contribution scores of gene $j$ with respect to subtype $p$

$gene\_scores\leftarrow\emptyset$;

for each matrix $scores$ in $(att\_scores, \bar{att\_scores} ):$

$raw\_scores\leftarrow mean\left( scores,axis=0 \right)$;

$raw\_scores\leftarrow mean\left( raw\_score_{j},axis=1 | all gene j have same gene name \right)$;

$raw\_scores$ $\leftarrow\{raw\_score_{j} |raw\_score_{j} >0)\}$

$gene\_scores.append(raw\_scores)$

${candidate\_genes}_{p}\leftarrow top\_k\left( gene\_scores,k \right)$;

end

return $candidate\_genes$

### 2. Supplementary Results

Due to the imbalance of the BRCA dataset, we have experimented several class-weighting options for the cross-entropy loss function. These options are unweighted, balanced, log-scale balanced, and manual design (**Figure S2**). Clearly, the forth strategy implementing the manual adjusment of the weight of each subtype shows considerably better performance on the classification capability comparing with the others. Specifically, the manually adjusted weights of 1, 1, 1, 5, and 5 corresponding the five labels: Basal, Her2, LumA, LumB, and Normal, respectively.

### 3. Supplementary Figures

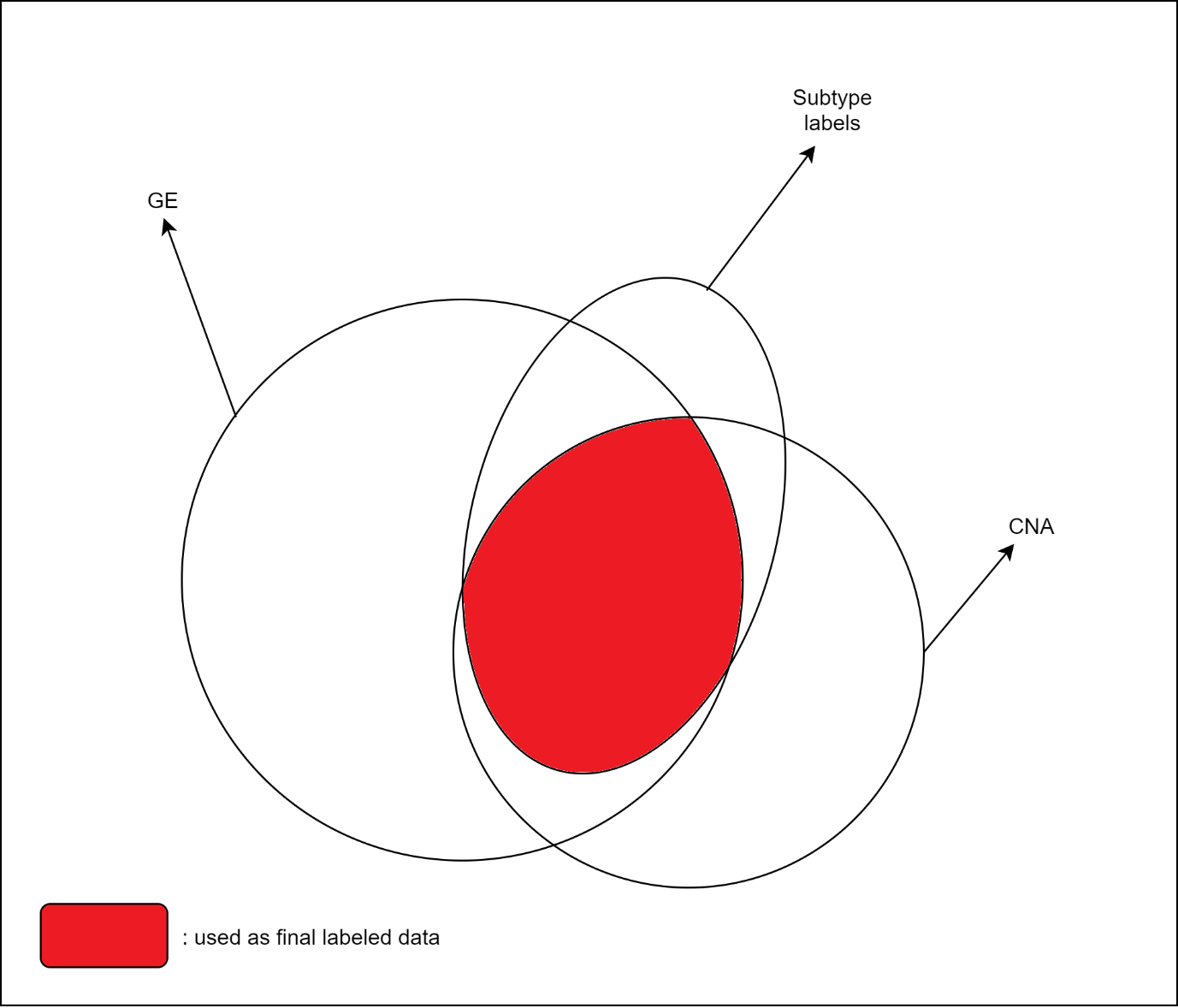

**Figure S1 |** **Illustration of the unlabeled and labeled data.** Red area in the oval ‘Subtype labels’ is the labeled data indicating that patients assigned to a specific subtype in both the GE data and CNA data. White areas in two circles ‘GE‘ and ‘CNA’ are the unlabeled data indicating that patients assigned to a specific subtype in only either of the GE data or CNA data, or not assigned to any subtype in both GE data and CNA data.

a

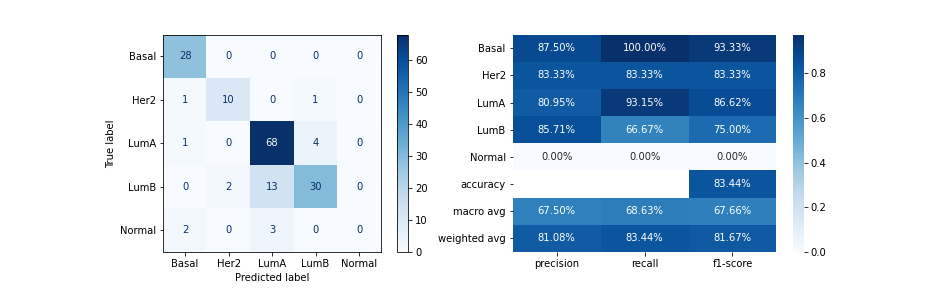

b

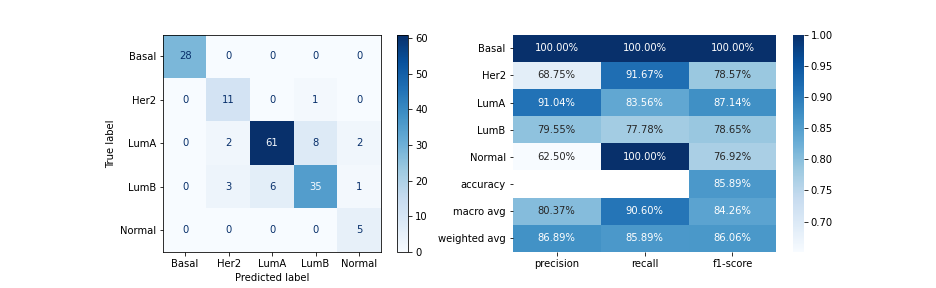

c

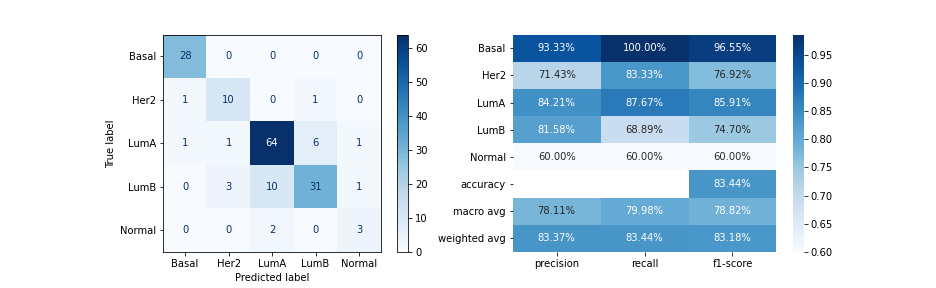

d

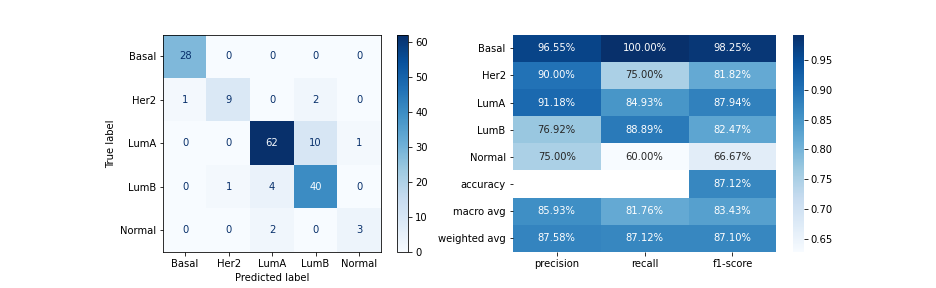

**Figure S2 |** **Classification performance comparisons in terms of different weighting strategies.** Our model is trained with different cross-entropy loss function to assess impact of each weighting strategy. **a** Unweighted. **b** Balance. **c** Log-scale balance. **d** Manual design. On the left hand side of each result is confusion matrices whose each row of the matrices represents the instances in an actual class and each column represents the instances in a predicted class. On the other side is the classification heatmaps, which summarize information of several metrics like precision, recall, f1 score of each class, macro average and weighted average of these metrics and overall accuracy. The best way to weigh the loss function is the manual design with the weights of 1, 1, 1, 5, and 5 correspond to the five labels Basal, Her2, LumA, LumB, and Normal-like, respectively.
